# BRD2 regulation of sigma-2 receptor expression upon cytosolic cholesterol deprivation

**DOI:** 10.1101/748236

**Authors:** Hongtao Shen, Jing Li, Xiujie Xie, Huan Yang, Mengxue Zhang, Bowen Wang, K. Craig Kent, Jorge Plutzky, Lian-Wang Guo

## Abstract

Traditionally a pharmacologic target for antipsychotic treatment, the sigma-2 receptor (S2R) was recently implicated in cholesterol homeostasis. Here we investigated the transcriptional regulation of S2R by the Bromo/ExtraTerminal epigenetic reader family (BETs, including BRD2, 3, 4) upon cholesterol perturbation.

Cytosolic cholesterol deprivation was induced using an export blocker of lysosomal cholesterol in ARPE19 cells. This condition upregulated mRNA and protein levels of S2R, and of SREBP2 but not SREBP1, transcription factors key to cholesterol/fatty acid metabolism. Silencing BRD2 but not BRD4 (though widely deemed as a master regulator) or BRD3 prevented S2R upregulation induced by cholesterol deprivation. Silencing SREBP2 but not SREBP1 diminished S2R expression. Furthermore, BRD2 co-immunoprecipitated with the SREBP2 transcription-active N-terminal domain, and chromatin immunoprecipitation-qPCR showed a BRD2 occupancy at the S2R gene promoter.

In summary, this study reveals a novel BRD2/SREBP2 cooperative regulation of S2R transcription in response to cytosolic cholesterol deprivation, thus shedding new light on epigenetic control of cholesterol biology.

## Introduction

After decades of studies, cholesterol biology remains inadequately understood, in particular, the regulations involving lysosomes, which distribute cholesterol to other organelles^1^. Recently, TMEM97 was reported as a novel player in cholesterol transport^2, 3^. An ER resident protein, TMEM97 can translocate to the lysosomal membrane where it appears to attenuate the activity of NPC1 (Niemann-Pick disease, type C)^3^, the transporter that “pumps” cholesterol out of the lysosome. TMEM97 was also suggested to interact with LDLR thereby involved in cholesterol uptake^3^. Intriguingly, the coding gene of the sigma-2 receptor (S2R), an enigmatic drug binding site pharmacologically identified 40 years ago, was finally (in 2017) unveiled to be TMEM97^4^.

There are a very small number of papers published on TMEM97 (a.k.a. MAC30)^5^; studies on S2R are largely limited to pharmacology such as anti-psychotic treatments^6^. As a result, little is known about the molecular regulations of S2R/ TMEM97/ MAC30 (hereafter denoted as S2R for clarity)^4^. S2R is highly expressed in progressive tumors and thereby targeted as a biomarker for diagnostic imaging^7^. Silencing S2R appeared to alleviate Niemann-Pick disease condition in a mouse model, which features mutated NPC1 and consequential cholesterol accumulation in lysosomes^2^. It is thus important to understand how S2R expression is controlled, whereas little is known at present.

Gene transcription programs are coordinately governed by transcription factors (TFs) and epigenetic factors. Rapidly growing literature supports a notion that the family of BETs (Bromo/ExtraTerminal-domain proteins) act as epigenetic determinants of transcription programs^8^. Among the BETs (BRD2, 3, 4), BRD4 is best studied and thought to function as a lynchpin-organizer of transcription assemblies. While its C-terminal domain (absent in BRD2 and BRD3) interacts with the transcription elongation factor (pTEFb) that activates the RNA polymerase II, its two bromodomains read (bind) acetylation bookmarks on histones and transcription factors. These interactions usher a transcription assembly to specific genomic loci^9^. BRD4 has been shown to play a master role in broad cellular processes ranging from proliferation to differentiation^10^ to autophagy^11^. Curiously, whether BETs are directly involved in cholesterol homeostasis remained unclear^12^. Relevant to this question, recent studies identified a crucial role of BRD4 in adipogenesis^10^. We were thus encouraged to explore whether BRD4 regulates the expression of S2R, a novel modulator of cholesterol transport^2^.

Herein we found that pan-BETs inhibition abolished S2R upregulation that was induced by cytosolic cholesterol deprivation. However, it was BRD2 but not BRD4 or BRD3 that was responsible for this BET function. This was unexpected given that BRD4 has been widely reported to be the determinant BET in diverse processes^8^. We also found that BETs inhibition repressed the transcription of both SREBP1 and SREBP2, the master TFs governing fatty acid and cholesterol homeostasis, and silencing SREBP2 but not SREBP1 inhibited S2R expression. Further results suggested a novel epigenetic mechanism whereby BRD2/SREBP2 co-occupy promoter regions of the S2R gene activating its transcription. Considering that BETs and S2R are targets of clinical (or trial) drugs^6, 13^, our findings may impact the studies on epigenetic regulations of cholesterol biology as well as translational medicine.

## Results

### Pan-BETs inhibition represses S2R expression that is stimulated by cytosolic cholesterol deprivation

To study epigenetic regulations of S2R, we chose ARPE19, a human epithelial cell line where cholesterol plays a critical role in cellular function/dysfunction^14^. To establish a cellular model to monitor S2R level changes, we first tested commonly used cytokine stimulants, including PDGF-AA, PDGF-BB, and TGFβ1. However, they did not significantly alter S2R mRNA levels (Figure S1). We then tested U188666A (hereafter abbreviated as U18), an NPC1 inhibitor that keeps cholesterol trapped inside the lysosome thereby generating a cholesterol-deprived cytosolic environment. We found that treatment with U18 increased S2R mRNA by up to 8 fold and S2R protein by ~ 2 fold. Interestingly, pretreatment with JQ1 (first-in-class BETs-selective inhibitor)^15, 16^ abrogated this U18-induced S2R upregulation (Figure 1, A and C). In contrast to U18 treatment, serum starvation had a relatively minor effect on S2R mRNA and protein upregulation (Figure 1, B and D). Confirming the function of U18 in reducing cytosolic cholesterol, Figure 1E showed that filipin staining of cholesterol was largely absent in the cytosol in U18-treated ARPE19 cells. The staining instead accumulated in perinuclear structures, which are typically known to be lysosomes^3^. By contrast, decrease of cytosolic cholesterol was less prominent under the starvation condition, consistent with a minor effect on S2R upregulation.

**Figure 1.**
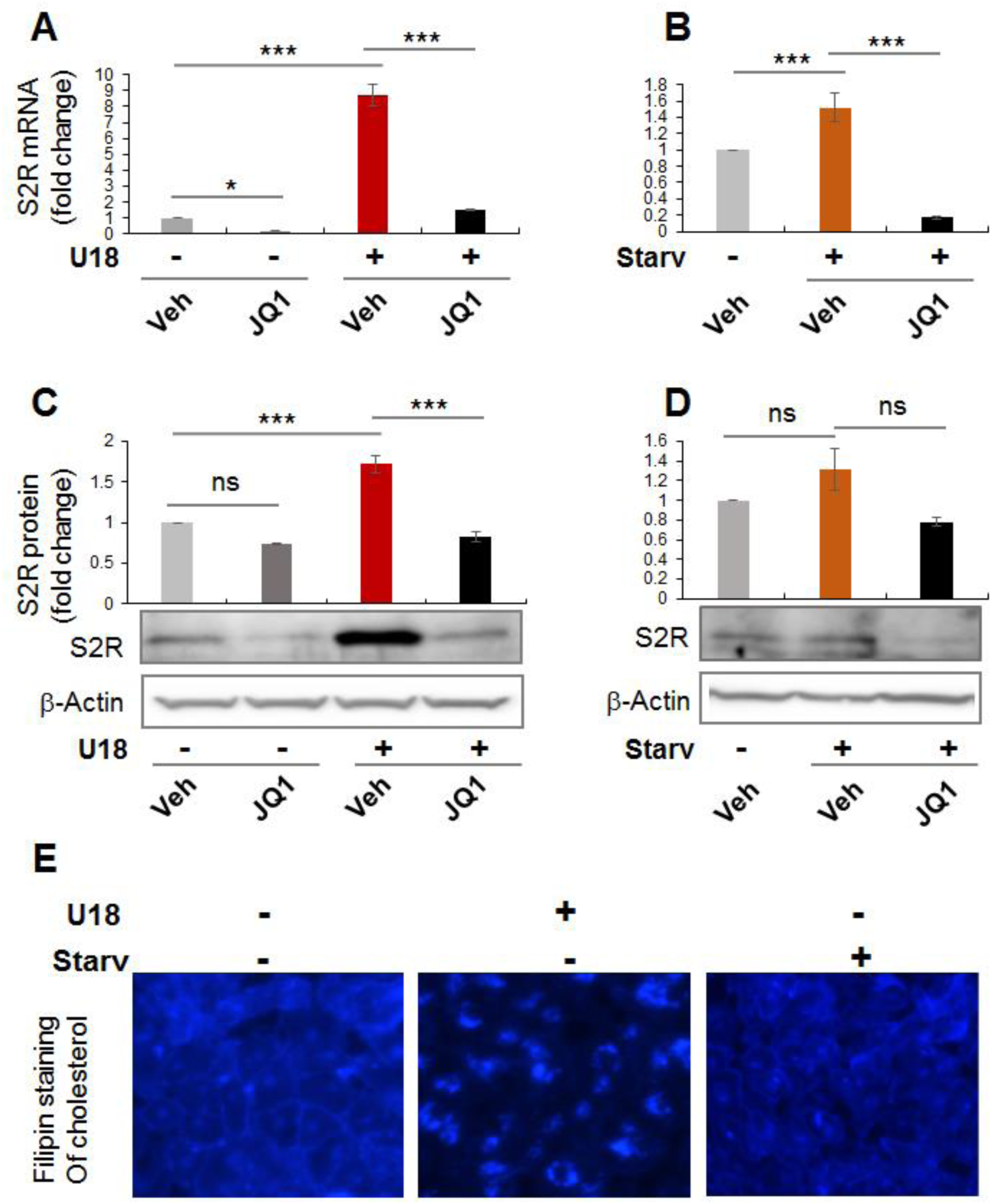
Pan-BETs inhibition prevents S2R upregulation upon cytosolic cholesterol deprivation. ARPE19 cells were cultured to an ~70-80% confluency in the DMEM/F12 medium containing 10% FBS. The cells were incubated with U188866A (abbreviated as U18, final 5 μM) for 24h before qRT-PCR (A and B) assay, Western blotting (C and D), or filipin staining of cholesterol (E). For starvation treatment, the medium was changed to that containing 0% FBS. For pretreatment to inhibit BETs, JQ1 (1 μM) or vehicle control (equal amount of DMSO) was included in the cell culture during the treatment with U18 or starvation. Quantification: At least three independent repeat experiments were performed; data were normalized to GAPDH (qRT-PCR) or β-actin (Western blot) and then to the control of vehicle only (no U18, no starvation). The normalized data were averaged (n ≥3) to calculate mean ± SEM. Statistics: One-way ANOVA with Bonferroni post-hoc test; *P<0.05, ***P<0.001; ns, no significance.

Taken together, whereas cytosolic cholesterol deprivation dramatically elevated S2R expression, pan-BETs inhibition with JQ1 abolished this upregulation.

### Silencing BRD2 but not BRD4 or BRD3 reduces S2R mRNA and protein

JQ1 is highly selective to the BET family yet it is a pan inhibitor that blocks the bromodomains in all BETs^16^. We therefore next determined by gene silencing which BET mainly accounted for the U18-induced S2R upregulation. Since there was no previous report to follow for silencing BETs in ARPE19 cells, we first tried lentiviral expression of shRNAs. BRD3 and BRD4 were effectively silenced by their respective shRNAs (Figure 2, A-C), yet BRD2 silencing was inefficient and we therefore used BRD2 siRNA instead. Given a wealth of literature evidence indicating BRD4 as a powerful regulator in a broad range of processes^17^, we expected BRD4 to be the determinant BET. To our surprise, silencing neither BRD4 nor BRD3 reduced U18-induced S2R expression (Figure 2, D-F); rather, silencing BRD2 attenuated U18-induced S2R (mRNA and protein) upregulation. To solidify this conclusion, we used two BRD3 shRNA sequences and two BRD4 shRNA sequences (Figure 2, B and C) (Figure S2) and furthermore, a BRD4 siRNA (Figure 3), but neither reduced S2R expression.

**Figure 2.**
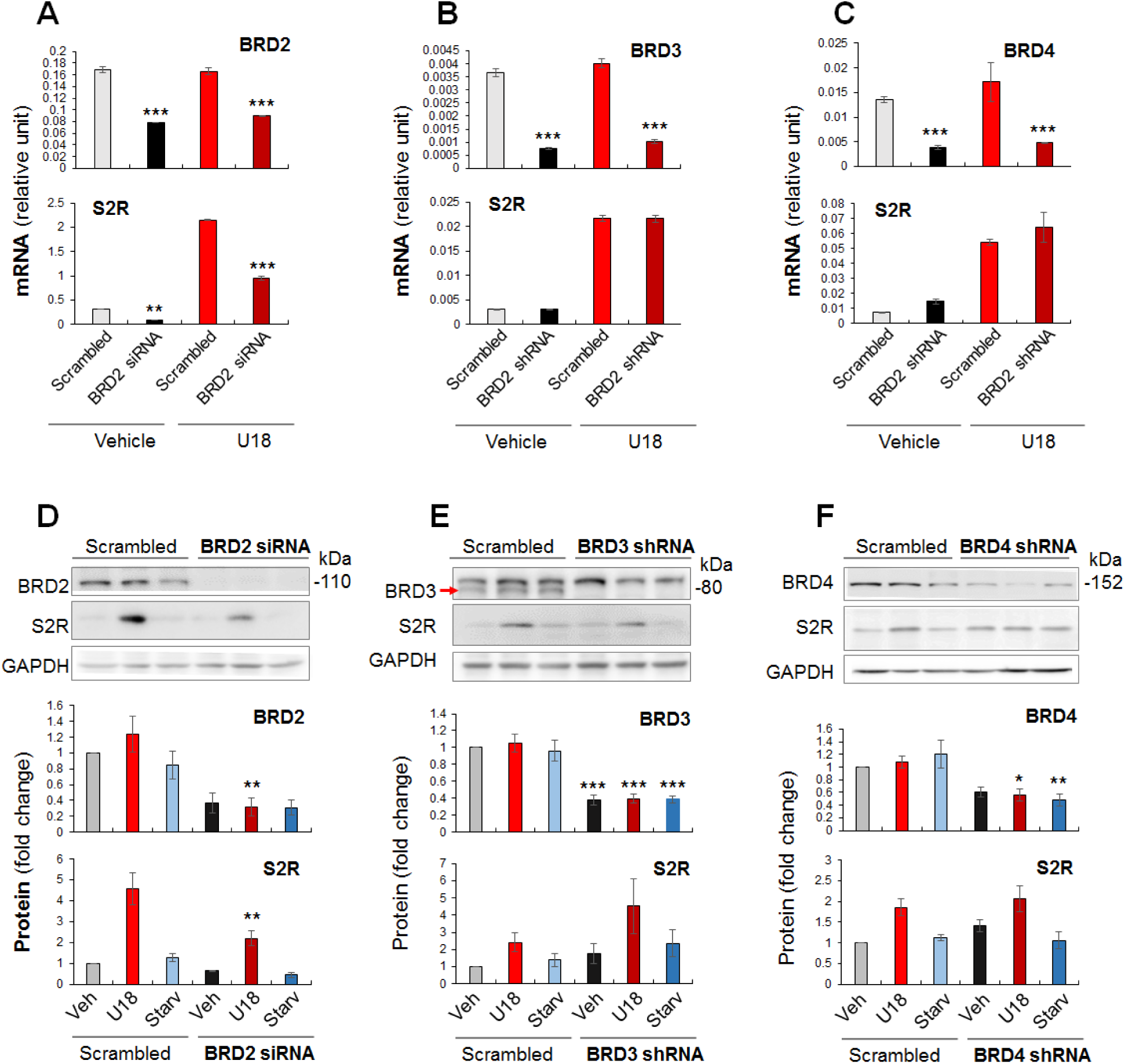
Inhibitory effect of BRD2 silencing on S2R mRNA and protein expression. ARPE19 cells were transfected with a BRD2-specific siRNA or infected with lentivirus to express a BRD3- or BRD4-specific shRNA prior to a 24-h U18 or starvation treatment, as described in detail in Methods. Cells were then harvested for assays. A-C. qRT-PCR. Shown is each representative of two similar yet independent experiments. Mean ± SD, n =3. D-F. Western blot. Data were quantified as described in the Figure 1 legend. Mean ± SEM, n =4 independent repeat experiments. Statistics (A-F): One-way ANOVA with Bonferroni post-hoc test; *P<0.05, **P<0.01, ***P<0.001, compared between conditions represented by light- and dark-colored bars (e.g. scrambled and BRD2 siRNA represented by light red and dark red in A, respectively); for simplicity, non-significant comparison is not labeled.

**Figure 3.**
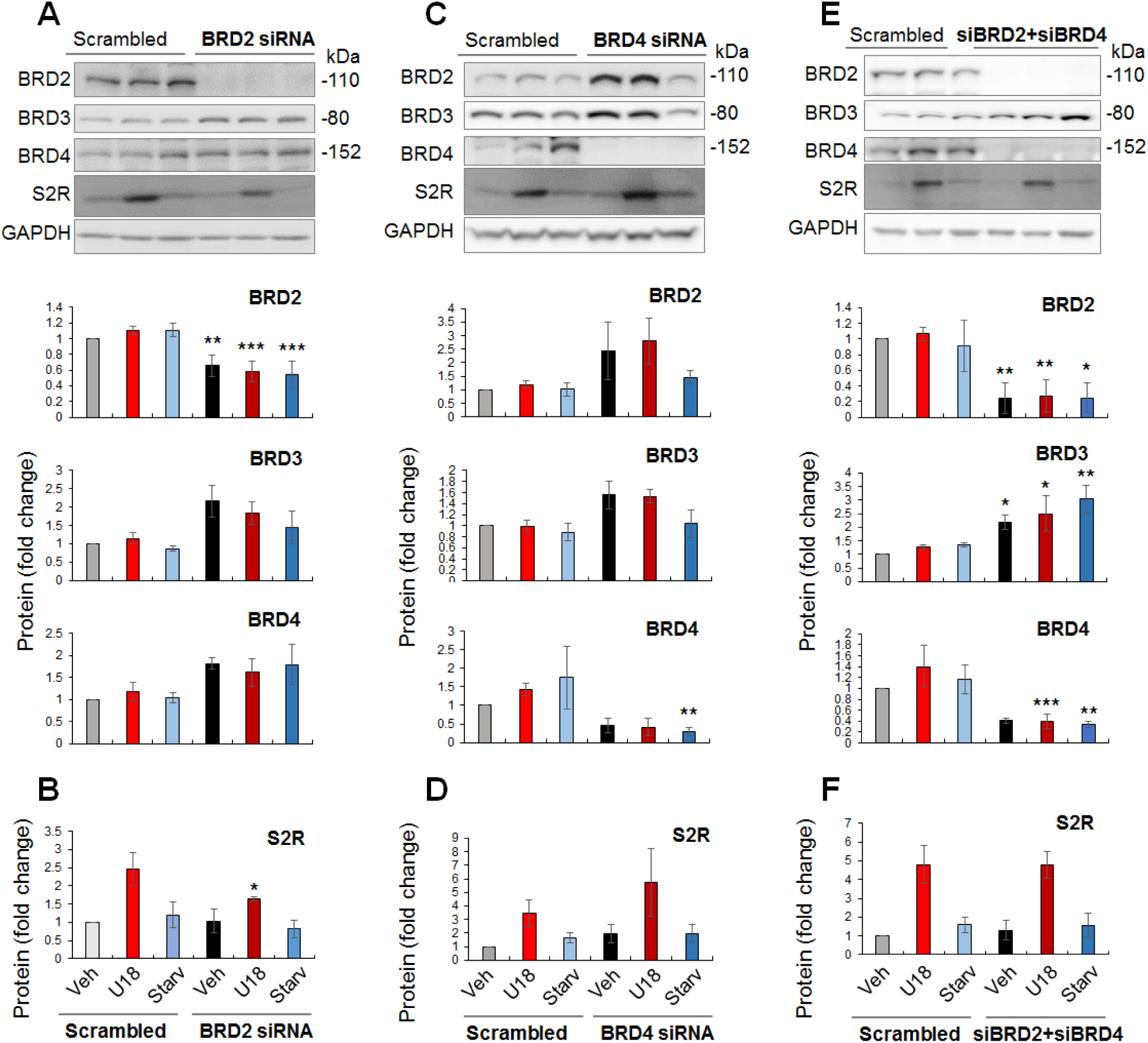
Effect of BRD2/BRD4 double silencing on S2R protein levels. Experiments were performed as described for Figure 2 except that BRD2 and BRD4 were silenced either separately with their specific siRNAs or simultaneously with combined siRNAs. A (BETs) and B (S2R) were from the same set of experiments, and so were C/D and E/F. Data were quantified as described in the Figure 1 legend. Statistics: One-way ANOVA with Bonferroni post-hoc test; n =4 independent repeat experiments; *P<0.05, **P<0.01, ***P<0.001, compared between conditions represented by a pair of light- and dark-colored bars (e.g. grey and black: scrambled and BRD-specific siRNA); non-significant comparison is not labeled.

It is noteworthy that BRD4 knockdown appeared to increase BRD2 protein and vice versa though without reaching statistical significance (Figure 3, A-D). An interesting question thus arose as to whether BRD4 knockdown could have raised S2R levels by increasing BRD2. We then silenced BRD4 in addition to BRD2 (Figure 3E), and the decrease of S2R became no longer evident (Figure 3F). The simplest explanation is that the changes of S2R toward opposite directions caused by silencing BRD2 and BRD4 canceled each other. Therefore, this result further supports a role for BRD2 in positively regulating S2R expression.

### Silencing SREBP2 but not SREBP1 represses S2R mRNA and protein expression

In a previously reported RNAi screening study, S2R (TMEM97) was found to be a target gene of the transcription factor SREBP2^3^. However, detailed studies on S2R transcriptional activation were not available, and whether S2R is under the control of the functionally paired transcription factor SREBP1 was not known. We thus performed silencing of SREBP2 and SREBP1 to determine an effect on S2R expression, either under U18 or starvation treatment. As shown in Figure 4A, whereas S2R mRNA levels were elevated by either U18 or starvation (to a lesser extent), SREBP2 silencing diminished this upregulation. By contrast, SREBP1 silencing did not reduce but rather, further enhanced S2R mRNA expression (Figure 4B); this enhancement was paralleled by increased BRD2 mRNA levels. Similar results occurred at the protein level; i.e. SREBP2 silencing abrogated S2R protein upregulation without changing BRD2 protein levels (Figure 4C), however, SREBP1 silencing appeared to increase BRD2 and S2R protein (no significance though) (Figure 4D). Of note, silencing SREBP2 reduced S2R expression without altering BRD2 mRNA or protein levels (Figure 4C), suggesting that SREBP2 is possibly downstream but not upstream of BRD2.

**Figure 4.**
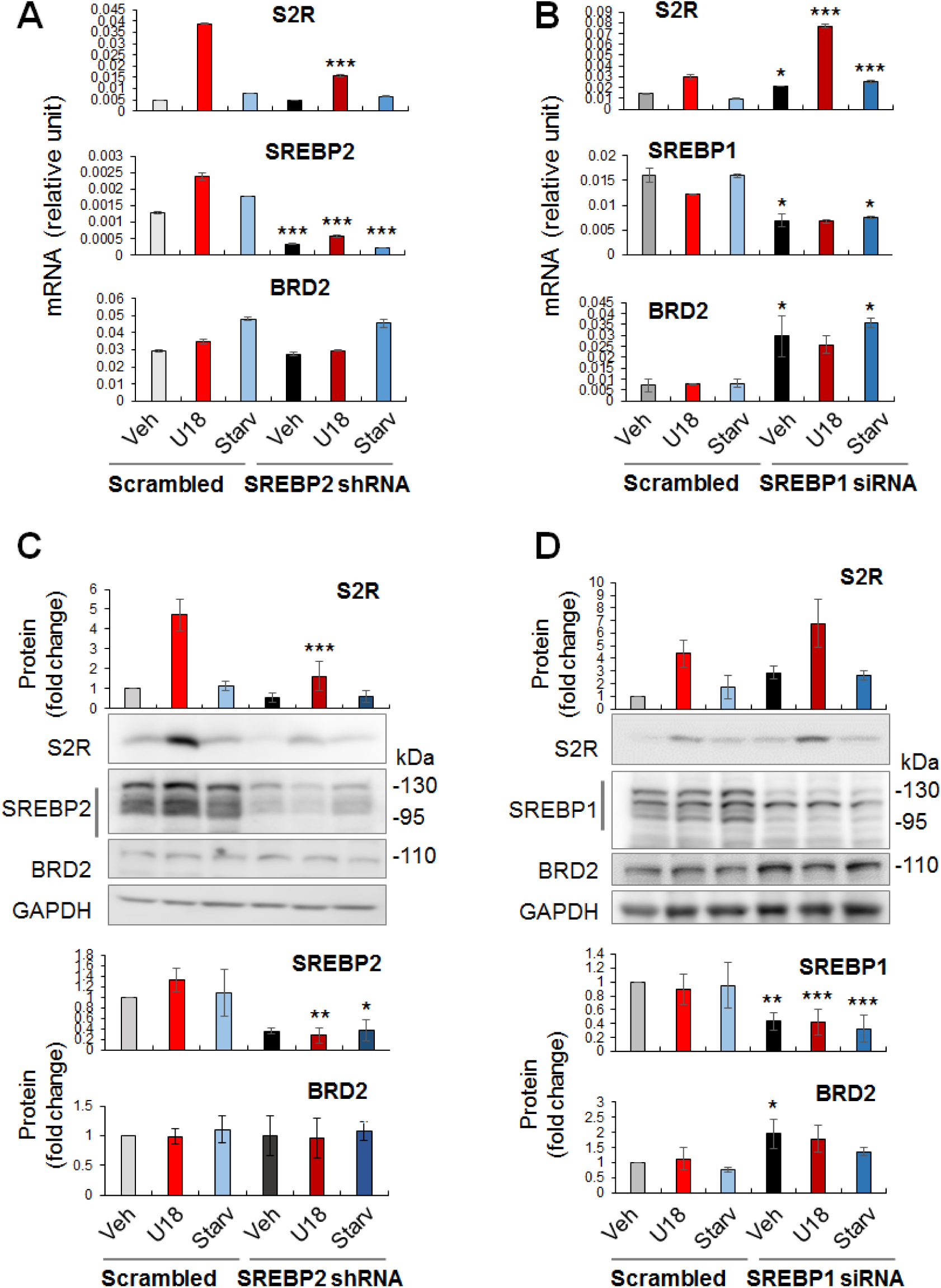
Effect of SREBP2 or SREBP1 silencing on S2R expression. Experiments were performed as described for Figure 2 except that SREBP2 or SREBP1 silencing was induced. A and B. qRT-PCR. Each represents one of two similar experiments. C and D. Western blot. Data were quantified as described in the Figure 1 legend. Statistics: One-way ANOVA with Bonferroni post-hoc test; n =4 independent repeat experiments (C and D); *P<0.05, **P<0.01, ***P<0.001, compared between conditions represented by a pair of light- and dark-colored bars (e.g. red and dark red); for simplicity, non-significant comparison is not labeled.

### Pan-BETs inhibition suppresses SREBP2 and SREBP1 mRNA levels

Given that S2R expression was controlled by both BRD2 and SREBP2, an epigenetic factor and a transcription factor, respectively (Figures 1–4), we next asked whether BRD2 regulated S2R expression via SREBP2. As shown in Figure 5 (A-D), treatment with U18 increased SREBP2 mRNA (by ~4 fold) and protein (albeit statistically insignificant), and SREBP1 mRNA to a minor extent. In contrast to U18, the starvation condition upregulated both SREBP2 and SREBP1 mRNA (by 10-15 fold) and protein levels (Figure 5, E-H). Remarkably, pretreatment with JQ1 abolished all of these changes. Moreover, using a different cell line (HEK293), we also observed upregulation of SREBP2 and S2R in U18-treated cells and its abolishment by pretreatment with JQ1 (Figure 5, I-K). These results suggest that the BETs family control the transcription of SREBP2 and SREBP1, two key TFs in cholesterol and fatty acid metabolism. To the best of our knowledge, BETs epigenetic regulations of SREBPs have not been previously determined in the specific context of cholesterol perturbation.

**Figure 5.**
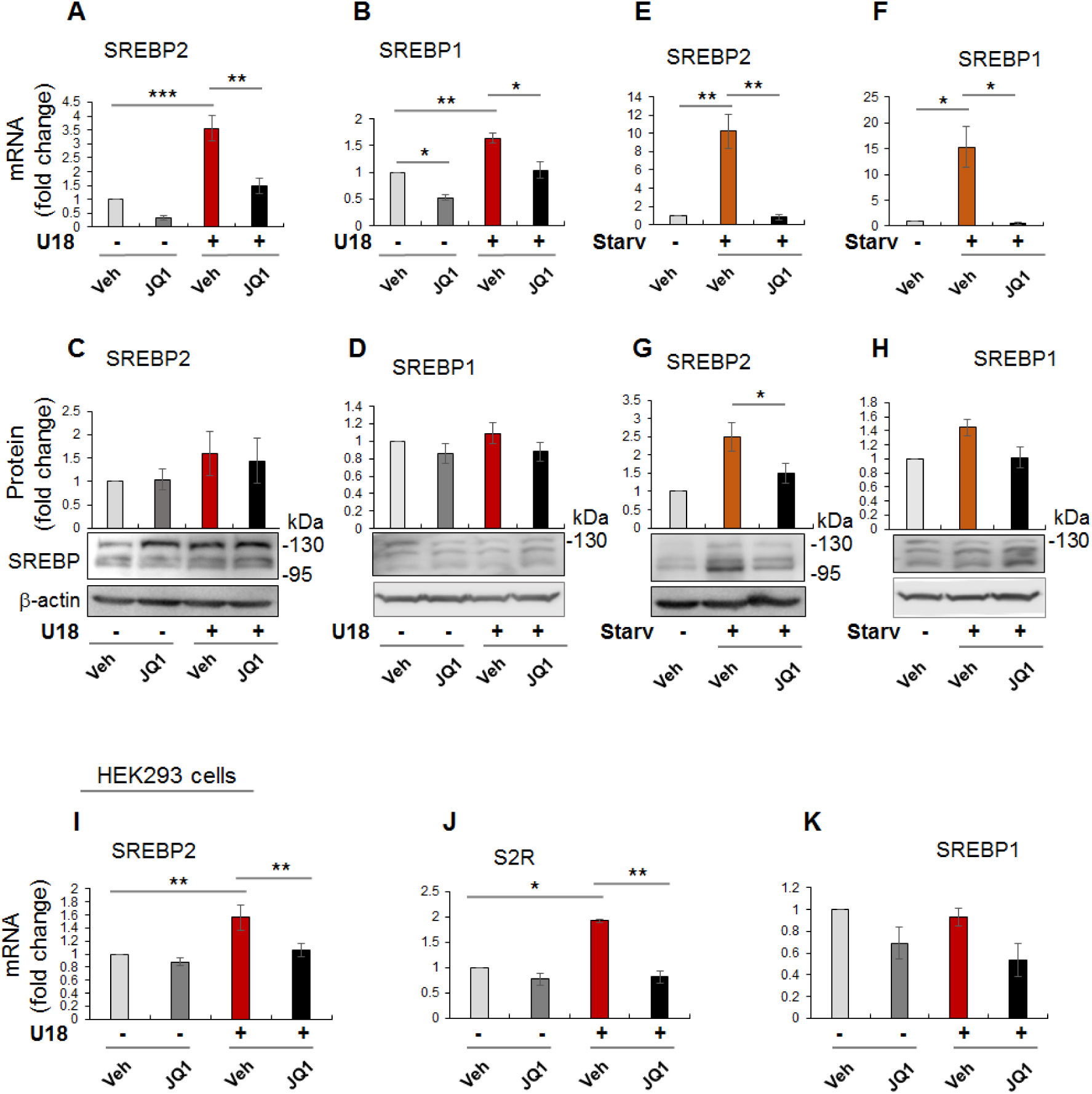
Pan-BETs inhibition suppresses the transcription of SREBPs. Experimental procedures and data quantification methods were the same as that described for Figure 1 (different sets of experiments) except that the main readouts were SREBP2 and SREBP1 mRNA (A, B, E, F, I, J, K) and protein (C, D, G, H) levels. Statistics: One-way ANOVA with Bonferroni post-hoc test; n =3 (qRT-PCR) or 4 (Western blot) independent repeat experiments; *P<0.05, **P<0.01, ***P<0.001; for simplicity, non-significant comparison is not labeled.

We then determined whether silencing a BET influences SREBP protein production. Figure 6 shows that neither BRD2 nor BRD4 silencing significantly altered SREBP2 or SREBP1 protein levels. This lack of changes in SREBP protein levels may be rationalized by post-transcriptional regulations.

**Figure 6.**
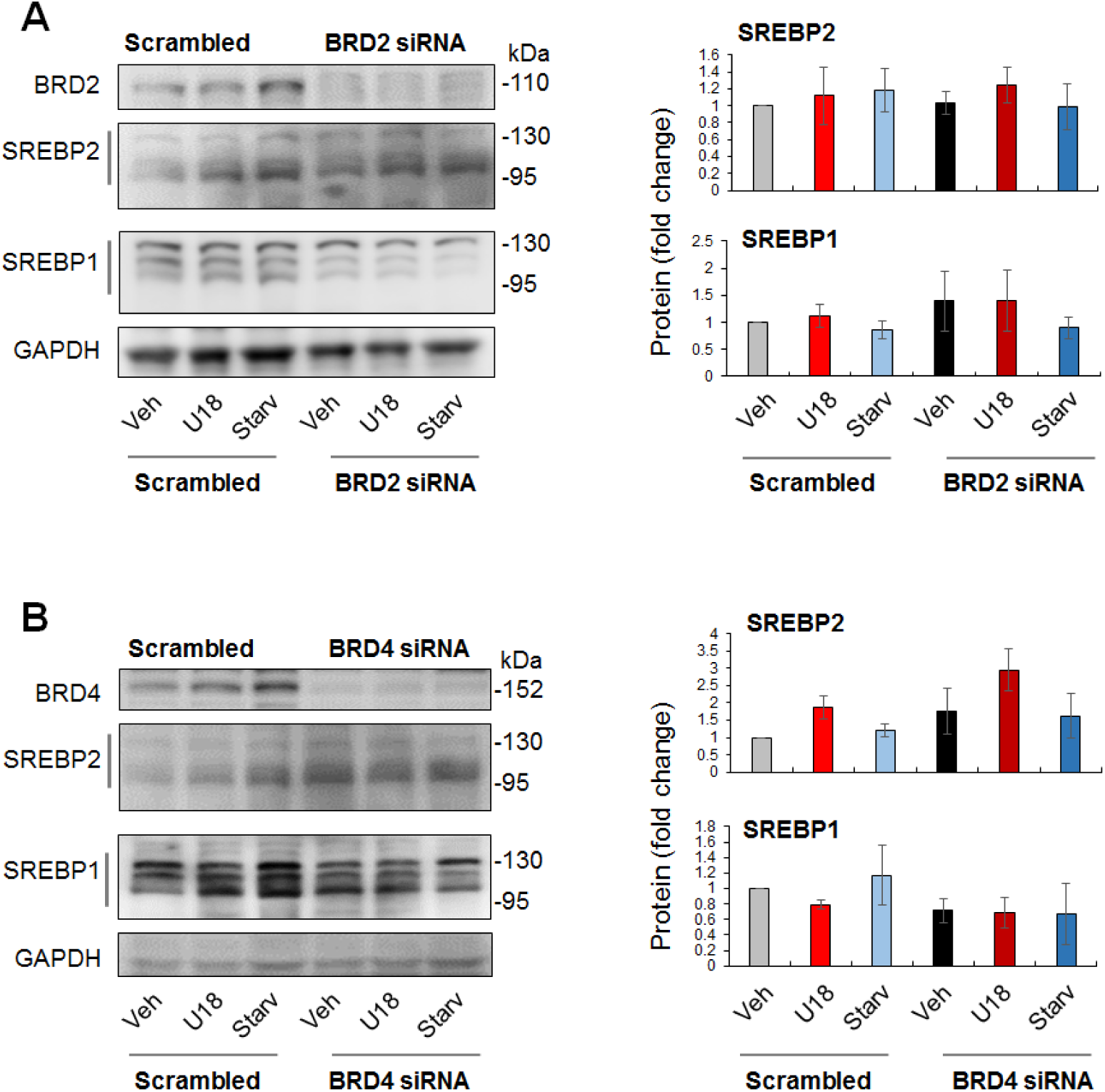
Effect of BRD2 or BRD4 silencing on SREBP2 or SREBP1 protein levels. Experimental procedures and data quantification methods were the same as described for Figure 2 (different sets of experiments), except that SREBP2 and SREBP1 protein levels were assessed as a result of BRD2 or BRD4 silencing. Statistics: One-way ANOVA with Bonferroni post-hoc test; n =4 independent repeat experiments; for simplicity, non-significant difference (between a pair of light and dark colors) is not labeled.

### BRD2 co-immunoprecipitates with the SREBP2 transcription-active N-terminal domain

Accumulating evidence suggests that BETs cooperate with specific TFs to assume transcriptional activation of select sets of genes, by two possible ways: altering the TF protein level or forming a complex with the TF to activate the transcription of target genes. Since the former did not occur (Figure 6), we next determined whether BRD2 forms a complex with the SREBP2 protein in the regulation of S2R transcription. Indeed, our data indicated that BRD2 co-immunoprecipitated with the SREBP2 N-terminal half molecule (Figure 7A), which is known to translocate into the nucleus to assume the SREBP2 TF function^3^. The specificity of this co-IP is manifested by the lack of BRD2 pulldown in the empty vector control (FLAG-GFP). The specific BRD2/SREBP2(N-term) co-IP was observed either without (Figure 7A) or with (Figure S3) U18 treatment.

**Figure 7.**
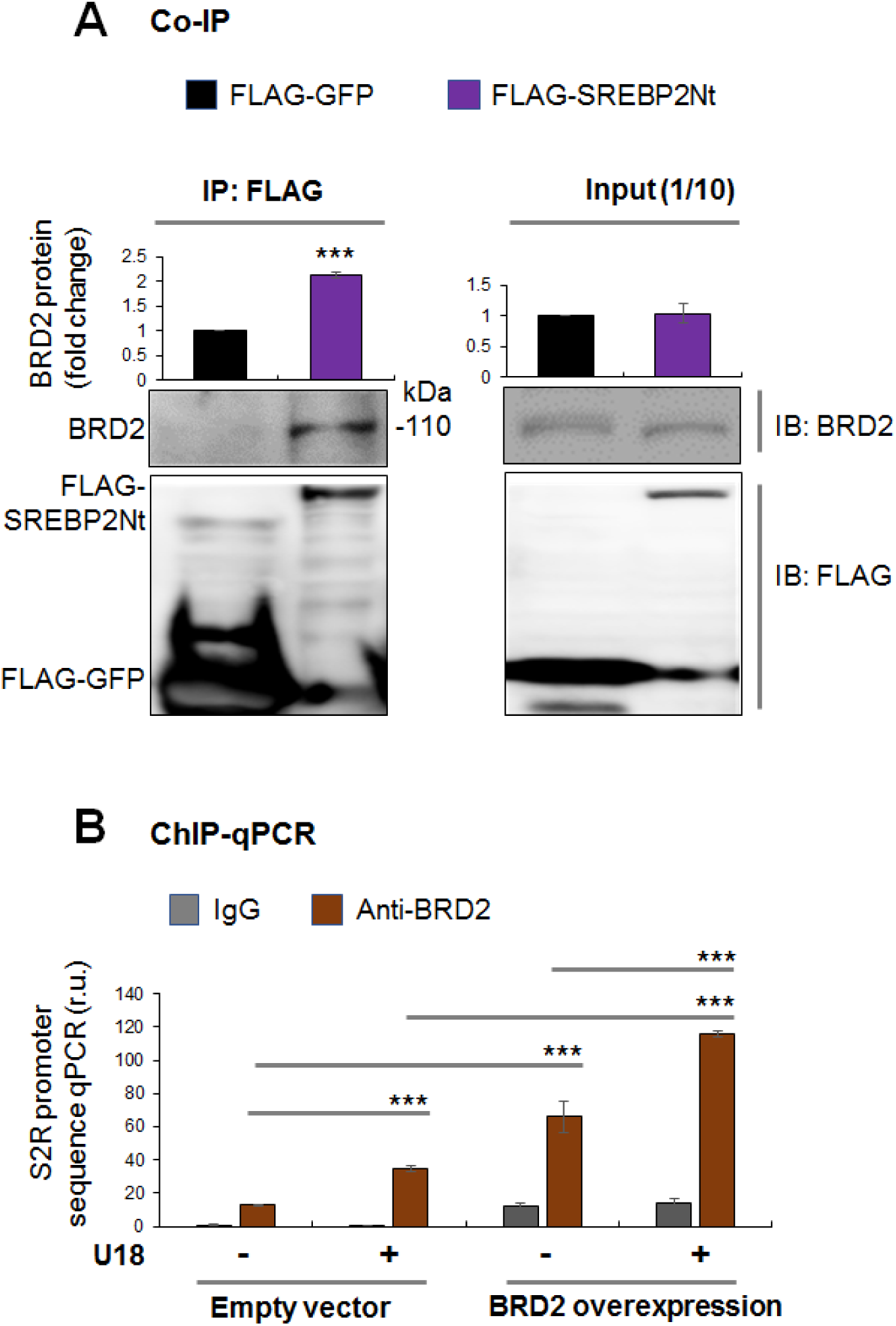
BRD2 co-immunoprecipitates with the SREBP2 N-terminal half molecule and S2R gene promoter regions. Co-immuoprecipitation (A) and ChIP-qPCR (B) experiments were performed as described in detail in Methods. **A.** Co-immuoprecipitation. Cells were transfected with a vector expressing FLAG-GFP (empty vector control) or FLAG-SREBP2Nt (N-terminal transcriptionally active). IP was performed with an anti-FLAG antibody. Presented blots represent one of three similar experiments (see Figure S3). Statistics: two-tailed paired Student t-test; ***P<0.001; n =3 independent repeat experiments. **B.** ChIP-qPCR. Cells were transfected with an empty vector or BRD2-expressing vector. ChIP was performed using an anti-BRD2 antibody. Presented is one of three similar experiments to detect different SREBP-binding regions (software-predicted) of the S2R gene promoter (see Figure S4). Mean ±SD; n =3 repeats. Statistics: One-way ANOVA with Bonferroni post-hoc test; ***P<0.001.

### BRD2 immunoprecipitates with SREBP-binding DNA regions of the S2R gene promoter

Finally, we used a BRD2 antibody for IP and performed ChIP-qPCR to detect S2R promoter regions that contain predicted SREBP-binding motifs (sterol response consensus elements)^3^. We found that while U18 treatment increased qPCR signal of a S2R promoter region by ~ 2 fold, overexpression of BRD2 further augmented the signal (Figure 7B). Similar results were obtained from experiments detecting other two different S2R gene promoter regions (Figure S4). The specificity of the ChIP-qPCR assay was confirmed by an outstanding signal-to-background ratio (~6-10 fold), i.e. a large difference in qPCR reading between the experiment using a BRD2 antibody for ChIP and that using IgG for control.

In aggregate, these results suggest that BRD2 upregulates S2R expression not by increasing the SREBP2 protein, rather, by forming a BRD2/SREBP2 complex that occupies S2R gene promoter regions to activate its transcription.

## Discussion

Cholesterol dysregulation leads to a myriad of pathological conditions^14^; S2R was recently suggested as a novel player in cholesterol intracellular transport^3^. To the best of our knowledge, the current report is the first to reveal a BETs-governed epigenetic control of S2R expression. We differentiated that silencing BRD2 but not BRD4 (though widely deemed as a master BET) or BRD3 effectively reduced S2R expression. While pan-BETs inhibition blocked the transcription of both SREBPs, silencing SREBP2 but not SREBP1 repressed S2R expression. Furthermore, our data provided evidence for that BRD2 controls S2R transcription not by increasing the SREBP2 protein but by forming a BRD2/SREBP2 complex at S2R gene promoter regions. Thus, our study suggests a previously unrecognized epigenetic mechanism whereby the duo of BRD2/SREBP2 positively regulate S2R expression in response to cholesterol deprivation in the cytosol.

The finding of BETs-dominated regulation of S2R expression is significant for the following reasons. (1) Epigenetics is crucial in cellular responses to extra- and intra-cellular environmental cues; however, little is known about epigenetic regulations of S2R^18^, whose expression is highly sensitive to cholesterol level perturbation. This knowledge gap likely stems from the fact that S2R is one of very few drug targets whose coding gene remained unknown until very recently^4^. (2) S2R was implicated (via pharmacology) in neurological diseases^19^ (e.g. Alzheimer’s)^20^, psychiatric disorders^6, 21^, and cancers^22^. In fact, S2R ligands have long been clinically used as antidepressants (e.g. haloperidol)^6^. Moreover, high S2R abundance was found in tumor tissues and cells, as detected with labeled S2R ligands^7, 22^ or unknowingly as TMEM97/ MAC30^23^. As such, S2R is often targeted for cancer imaging (e.g. PET scanning)^7^. More recently, it was reported that S2R (TMEM97) knockdown attenuated Niemann-Pick disease phenotypes in a mouse model, linking S2R to lysosomal cholesterol export^2^. (3) S2R was found to re-locate to lysosomes when intracellular cholesterol levels dropped^3^. It was thus speculated that S2R may interact with NPC1 modulating its cholesterol-exporting function in the lysosomal membrane, whereas with no direct evidence available. (4) Recent studies suggested that S2R co-localizes with LDLR, which via internalization carries esterified cholesterol into the cell^3^; S2R knockdown or knockout impairs cholesterol (and LDLR) uptake. Together, these studies indicated the biological importance of S2R and its regulators and motivated our investigation into BETs-dictated epigenetic regulation of S2R expression.

In this novel regulation, it is somewhat surprising that BRD2 rather than BRD4 is found to be the determinant BET. In contrast to BRD2 and BRD3, BRD4 has been intensively studied and shown to play a critical role in many crucial cellular processes and pathological conditions^8, 24^. The BRD4 molecule (*vs* BRD2 and BRD3) has nearly doubled length which contains a unique C-terminal domain. From a functional/structural perspective, BRD4 is dubbed a “Swiss army knife”^25^. Its two bromodomains “dock” the BRD4-organized regulatory complex (including TFs and cis-regulators) to specific bookmarked chromatin sites, with its C-terminal domain promoting transcriptional activation by interacting with the transcription elongation factor that in turn activates the RNA polymerase II. BRD4 was very recently found to possess intrinsic kinase^26^ and acetyl transferase activities^27^. However, in our specific experimental setting of U18-induced cytosolic cholesterol deprivation, it was BRD2 but not BRD4 that positively regulated S2R transcription. Inasmuch as BRD2 lacks a C-terminal domain and its bromodomain sequences are different from that of BRD4^16^, our results may implicate a BET mechanism distinct from that of BRD4. Given limited information about BRD2 functional mechanisms, future studies are needed to elucidate the molecular workings that underlie BRD2-dominated epigenetic control of S2R transcription.

To this end, the previous evidence for S2R being a target gene of SREBP2^3^ inspired our investigation that led to another novel finding, i.e. BETs govern the transcription of both SREBP1 and SREBP2, the TFs key to fatty acid/cholesterol regulations. This BETs control over SREBP’s transcription is exciting, given that BETs and SREBPs are both master regulators of vital cellular activities yet their relationship was previously unknown in the context of cholesterol homeostasis where SREBPs are critically important. Thanks to the recent discovery of inhibitors selective for BETs^15, 28^, important BETs functions have been recently identified. BETs inhibition was initially shown to be highly effective in blocking transcription programs of inflammation^28^ and oncogenic proliferation^29^. The importance of BET biology was then extended to stem cell differentiation, hematopoiesis, synaptic plasticity^12^, and recently, also adipogenesis^10, 30^. Most recently, BRD4 was found to regulate intra-nuclear/cellular processes as well, such as chromatin architectural remodeling^31^ and autophagy^11^. However, there is a dearth of information on a role for BETs in regulating SREBPs. The closest relevance is a report on BETs-associated super-enhancers formed during adipogenesis^10^. While SREBP1 (but not SREBP2) was on the list of JQ1-regulated genes derived from RNAseq data, the relationship between BETs and SREBP1 was not specifically examined. Increased HDL-cholesterol or decreased LDL-cholesterol was observed in animal plasma after treating with a BETs inhibitor^32, 33^. However, little is known about specific BETs-dominated cholesterol regulatory pathways, underscoring the importance of investigating BETs regulations of SREBPs and their downstream effector genes.

SREBPs could be reasonably categorized as master TFs. There are total over one thousand TFs, only a limited number of them are deemed master TFs. These master TFs often direct transcription programs that define a cell type or cell state^8^. As such, they are sensitive to extra- and intra-cellular environmental perturbations, and their expression levels and activities are key to cell state changes and associated disease conditions^34^. Recently, master TFs were found to co-localize with BRD4 in the genomic landscape thereby playing a critical role in inflammatory (e.g. NFkB)^34^, proliferative (c-Myc)^29, 35^, or immunological (T-bet) processes. Another prominent feature of master TFs is that they potently co-activate the transcription of specific sets of genes, by forming a complex with BETs to promote not only target gene expression but also their own transcription^8^. In consonance, herein our ChIP-qPCR data provided evidence for BRD2 occupancy at the SREBP2 gene promoter (Figure S5). Based on these criteria, SREBP2 and SREBP1 appear to be master TFs. SREBP1 and SREBP2 differentially regulate fatty acid and cholesterol pathways (though with possible crosstalk). This may explain our observation that SREBP2 but not SREBP1 positively regulated S2R expression upon cholesterol deprivation. Of note, serum starvation markedly stimulated the expression of both SREBP2 and SREBP1, likely because serum contains both cholesterol and fatty acids. Consistently, pan-BETs inhibition with JQ1 averted upregulation of both SREBP1 and SREBP2 mRNAs stimulated by serum starvation. In addition, JQ1 also reduced mRNAs of SREBP target genes tested herein including not only S2R, but NPC1, NPC2, and LXRs as well (Figure S6). As such, BETs together with SREBPs may govern transcription programs of the cholesterol /fatty acid pathways. For this possibility, future RNAseq studies are needed to provide comprehensive evidence.

Furthermore, our results suggest that in the condition of cholesterol deprivation the BRD2/SREBP2 duo accounted for the upregulation of S2R. Evidence includes the following: (1) U18 treatment dramatically upregulated S2R and SREBP2 but not SREBP1 expression. (2) Silencing SREBP2 but not SREBP1 reduced S2R mRNA and protein. (3) BRD2 co-immunoprecipitated with the SREBP2 N-terminal TF domain. (4) ChIP-qPCR assays using a BRD2 antibody suggested that BRD2/SREBP2 co-occupied S2R promoter regions, and this co-occupancy was enhanced by U18-induced cholesterol deprivation and BRD2 overexpression. This novel result is consistent with the well-established SREBP2 functional mechanism; i.e. SREBP2 in an inactive state resides in the endoplasmic reticulum (ER) membrane, but upon decrease of cytosolic (hence ER) cholesterol, SREBP2 is transported to Golgi and cleaved into two half molecules^3^. Whereas the C-terminal half stays in the cytosol, the N-terminal half enters the nucleus acting as a TF for the expression of genes involved in cholesterol metabolism and transport. Taken together, the pockets of new information obtained herein and that from the literature form a coherent picture of BRD2/SREBP2-dictated S2R transcription in response to cytosolic cholesterol perturbation.

## Conclusions

We present the first evidence on BETs control over the transcription of S2R, a recently unveiled player in cholesterol-associated regulations. Our results have sketched out a potential epigenetic regulatory mechanism whereby BRD2 forms a complex with SREBP2 that localizes at the S2R gene promoter activating S2R transcription. Further investigation may shed new lights on BRD2-dominated transcription programs that sensitively respond to cholesterol level changes. Along this line, studies on BETs and S2R, both targets of increasing clinical (trial) drugs^6, 13, 24, 36, 37^, may synergize interventional opportunities for cholesterol-associated pathological conditions.

## Acknowledgments and Sources of Funding

This work was supported by NIH R01 grants HL133665 and EY029809 to L.-W.G., HL143469 and HL129785 to K.C.K. and L.-W.G., and an AHA pre-doctoral award 17PRE33670865 (to M.X.Z.).

## Methods

### Major materials

JQ1 was purchased from Apexbio (A1910). U18666A was from Sigma-Aldrich (662015). Filipin complex was from Sigma-Aldrich (F9765). ARPE19 cells and HEK293 cells were obtained from American Type Culture Collection (ATCC). Scrambled and BRD2-, BRD4-, or SREBP1-specific siRNAs were from Thermo Fisher Scientific (scrambled: AM4635; BRD2: AM16708, ID-118266; BRD4: 4457298, ID-s23902; SREBP1: AM51331, ID-5140). Lipofectamine3000 was from Thermo Fisher Scientific (L3000008).

### Cell culture and treatment

ARPE19 cells were cultured in the DMEM/F12 medium (Thermo Fisher Scientific, 11320082) supplemented with 10% fetal bovine serum (FBS) and penicillin/streptomycin (Thermo Fisher Scientific, 5140163) at 37 °C in a humidified atmosphere with 5% CO_2_. HEK293 cells were maintained in DMEM (Thermo Fisher Scientific, 10569010) supplemented with 10% FBS and penicillin/streptomycin. To induce cytosolic cholesterol deprivation, ARPE19 cells were seeded in 6-well plates at 3×10^5^ cells/well and cultured for 24h, U18666A was then added (final 5 μM) and incubated for another 24h. In some experiments, JQ1 was added (5 μM) together with U18666A. For starvation experiments, the culture medium containing 10% FBS was changed to that without FBS in which cells were cultured for 24h. Cells were then collected for various analyses. For cholesterol staining, cells were washed 3x with PBS before fixing with 3% paraformaldehyde for 1h. The reaction was stopped with glycine (1.5mg/ml) and the cells were then stained for 2h at room temperature in the filipin working solution (0.05 mg/ml in PBS with 10%FBS). Images were taken with the Nikon fluorescence microscope using a UV filter set (340-380 nm excitation).

### RNA isolation, reverse transcription, and quantitative real-time PCR (qRT-PCR)

Total RNA was isolated and purified using Trizol Reagent (Thermo Fisher Scientific,15596026) following the manufacturer’s instruction. RNA was reverse-transcribed using the High-Capacity cDNA Reverse Transcription kit (Thermo Fisher Scientific, 4368814). cDNA of 1 μLfrom 20 μL reaction volume was amplified by real-time quantitative PCR (Applied Biosystems Quant Studio 3, Thermo Fisher Scientific) with Perfecta SYBR Green Master Fast Mix (VWR, 101414-286)^38^. Relative gene expression was determined by the 2^−ΔΔCt^ method, normalized to GAPDH, and presented as relative mRNA levels. qPCR analyses were done in triplicate. Experiments were repeated at least twice. Primers are listed in Table S1.

### Preparation of lentivectors for shRNA expression

The pLKO.1-puro empty vector was purchased from Addgene (#8453). A scrambled shRNA control and gene-specific shRNAs were designed through RNAi Central (http://cancan.cshl.edu/RNAi_central/step2.cgi). The corresponding oligonucleotides (ordered from Thermo Fisher Scientific) were annealed (95 °C to 25 °C, 0.1 °C /s) and cloned into the pLKO.1-puro vector, followed by confirmatory sequencing at the Ohio State University facility. The shRNA sequences (of final siRNA products) are listed in Table S2. For lentivirus packaging, lentivector plasmids were transfected into HEK293T cells together with packaging and envelope plasmids (psPAX2 and pMD2.G) using Lipofectamine 3000 (Thermo Fisher Scientific, L3000008). Three days after transfection, the medium was passed through a filter of 0.45 μm pore size and then used for transduction of ARPE19 cells. After 48 hours of infection, cells were selected with 1 μg/ml of Puromycin (Thermo Fisher Scientific, A1113803) for 5-10 days.

### Plasmid and siRNA transfection

Plasmids were transfected via co-incubation with Lipofectamine3000 for 24h in the recipient cell culture following the manufacturer’s instructions. The medium was then replaced with fresh DMEM/F12 for the cells to recover (24h) before their further use in various analyses.

For siRNA transfection, ARPE19 cells were cultured to 80% confluency in the DMEM/F12 medium containing 10% FBS in 6-well plates, and then added with a scrambled or BRD2-, BRD4-, or SREBP1-specific siRNA (sequence information available at the manufacturer, Thermo Fisher Scientific). The cells were transfected overnight using the Lipofectamine RNAi Max transfection reagent (Thermo Fisher Scientific, 13778150), and then recovered in the DMEM/F12 culture medium for 24h before further experimental use.

### Western blotting

Western blot analysis was performed following our published protocol^38^ with minor modification. Briefly, cells were lysed with the Pierce RIPA lysis buffer (Thermo Fisher Scientific, 89901) containing Halt Protease Inhibitor Cocktail (Thermo Fisher Scientific, 87785). Total protein concentration was determined using the DC Protein Assay kit (Bio-Rad, 5000111). The cell lysates were solubilized in Pierce Lane Marker Non-Reducing Sample Buffer (Thermo Fisher Scientific, 39001) and heated at 95°C for 10 min prior to SDS-PAGE and Western blotting. The information for the antibodies used are available in Table S3. Specific protein bands on Western blots were quantified by the ImageJ 64 software (https://imagej.nih.gov/ij/) using Gel analyzer script. Densitometry data were normalized to loading control (GAPDH or β-actin) and then to a basal condition (e.g. vehicle and/or scrambled sequence siRNA).

### Co-immunoprecipitation (co-IP)

pEGFP-N1-FLAG (empty vector) (Addgene, 60360) and pFLAG-SREBP2Nt (N-terminal transcriptionally active domain, amino acids 1-482) (Addgene, 26807) were used to transfect cells (HEK293). For co-IP, cells were lysed on ice for 30min in Pierce IP Lysis Buffer (Thermo Fisher Scientific, 87788) containing Halt Protease Inhibitor Cocktail (Thermo Fisher Scientific, 87785), and then centrifuged at 12000 rpm for 15min at 4°C. The supernatant was incubated with 50 μl of Pierce Anti-DYKDDDDK Magnetic Agarose beads (Thermo Fisher Scientific, A36797) at 4°C overnight. The beads were washed 3x with cold PBS buffer and then incubated in 0.1 M glycine (pH 2.8) for 10 min at room temperature with frequent vortex to elute the immuno-precipitates. The eluate was neutralized with 1M Tris-HCl, pH 8.5 (15 μl per 100 μl eluate), and briefly heated at 95°C prior to its use for SDS-PAGE and Western blot analysis.

### Chromatin immunoprecipitation (ChIP) analysis

ChIP analysis was performed by using the Pierce Magnetic ChIP kit (Thermo Fisher Scientific, 26157) and following the manufacturer’s manual. Briefly, formaldehyde (final concentration 1%) was incubated with the ARPE19 cell culture to cross-link protein with DNA for 10 min, and the reaction was then quenched with a glycine solution for 5 min. Cells were washed with ice-cold PBS, lysed in the Membrane Extraction buffer, and centrifuged at 3000g for 5 min to collect the nuclei. The nuclear pellets were re-suspended in 200 µl of MNase Digestion Buffer Working Solution and then digested by incubation with MNase at 37°C for 15 min. The reaction was terminated in MNase Stop Solution. The nuclei were recovered by centrifugation at 9000g for 5 min, re-suspended in IP Dilution Buffer, and then sonicated (four 5-second pulses at 20 Watts for 2X10^6^ cells) to break the nuclear membrane. Following centrifugation at 9000g for 5 min, the supernatant was collected and incubated overnight with a BRD2 antibody (Table S3) or IgG control (5 µg antibody per reaction). ChIP-grade Protein A/G Magnetic beads were added and incubated overnight at 4°C on mixing. The beads were collected and washed sequentially with IP Wash Buffer-1, IP Wash Buffer-2 and then resuspended in the elution buffer. The protein-DNA cross-link was reversed with 5 M NaCl followed by RNA and protein digestion with RNAse A and Proteinase K. The DNA pulldown was purified with DNA Clean-Up Column and used for qPCR. Primers used for detection of S2R and SREBP promoter regions that contain predicted SREBP-binding sterol-response consensus elements are listed in Table S4. For each qPCR assay, triplicate samples were used, and data were normalized to respective input samples. For BRD2 gain of function, we selected a BRD2-expressing stable ARPE19 cell line using a lentivector constructed based on an empty vector (Addgene, 19319).

### Statistical analysis

Repeat experiments were performed on different (n≥3) occasions. Results were plotted as mean ± SEM unless otherwise specified. Statistical significance (set at P<0.05) was determined by one-way ANOVA with Bonferroni post-hoc test for multi-group comparison or two-tailed paired Student’s t test for two-group comparison (GraphPad Prism 7). Significance is indicated as *P < 0.05, ***p < 0.01, or ***P <0.001; no significance is labeled as “ns” or not labeled, as specified in each figure legend.

---

**Figure S1.**
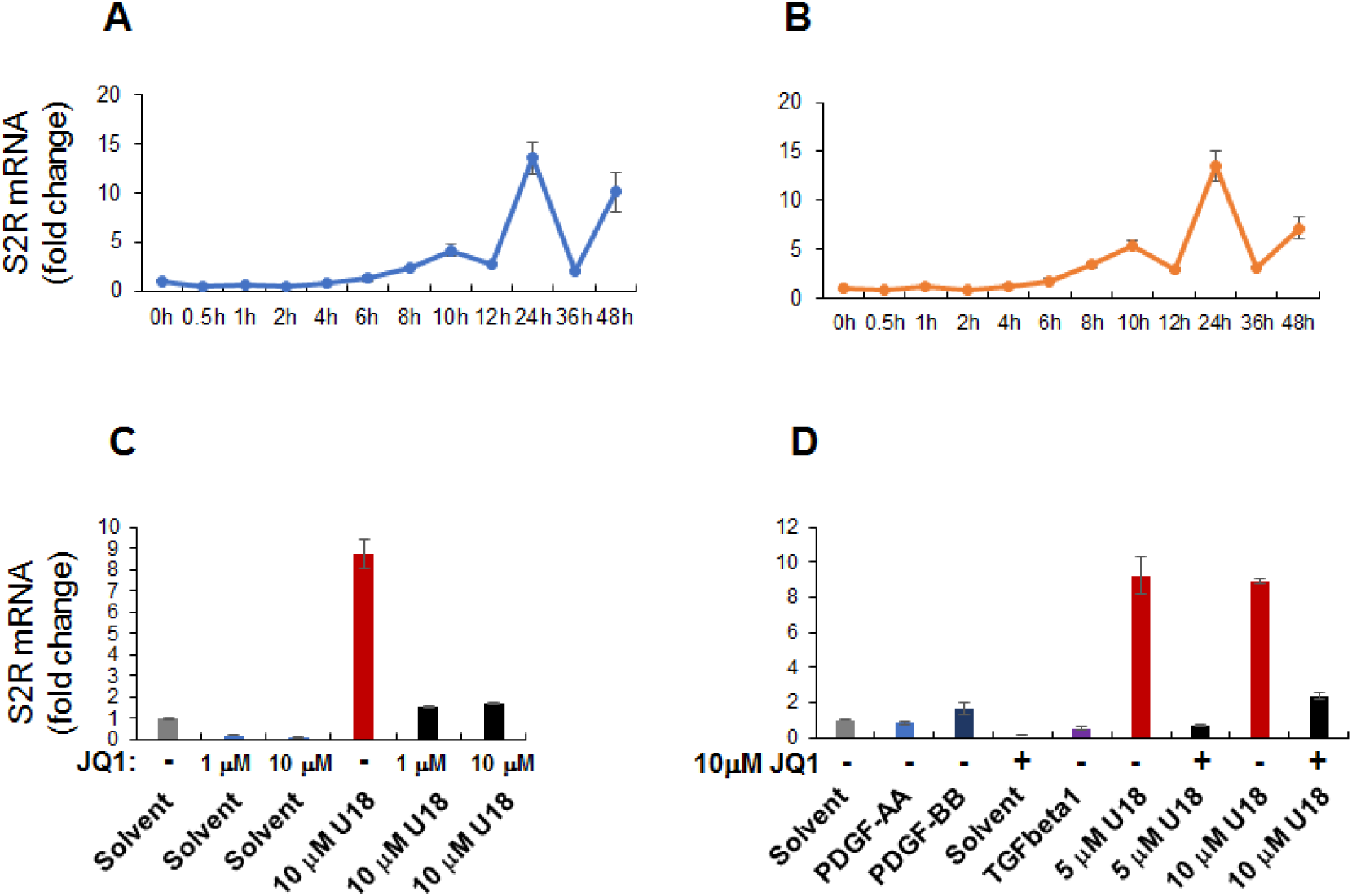
U18 induces upregulation of S2R mRNA in ARPE19 cells. Experiments were performed as described in Figure 1 except for the various conditions indicated in this figure. For each data point or bar, triplicate samples were used. A and B. Time course of U18 treatment of ARPE19 cells; two repeat experiments. The data indicates that S2R mRNA levels were maximally increased by U18 (5 μM) at 24h after treatment. C and D. Effects of cytokines and different concentrations of U18 and JQ1 on S2R mRNA expression. PDGF-AA, 20 ng/ml; PDGF-BB, 20 ng/ml; TGFβ1, 20 ng/ml. PDGF-AA (cat#1055AA050) and PDGF-BB (cat#520BB050) were from R&D Systems. Human recombinant TGFβ1 was from Thermo Fisher Scientific (cat#PHG9214). The data indicate that treatments with 5 μM and 10 μM U18 were equally effective in inducing S2R mRNA upregulation, and 1 μM JQ1 was as potent as 10 μM in blocking U18-induced S2R upregulation.

**Figure S2.**
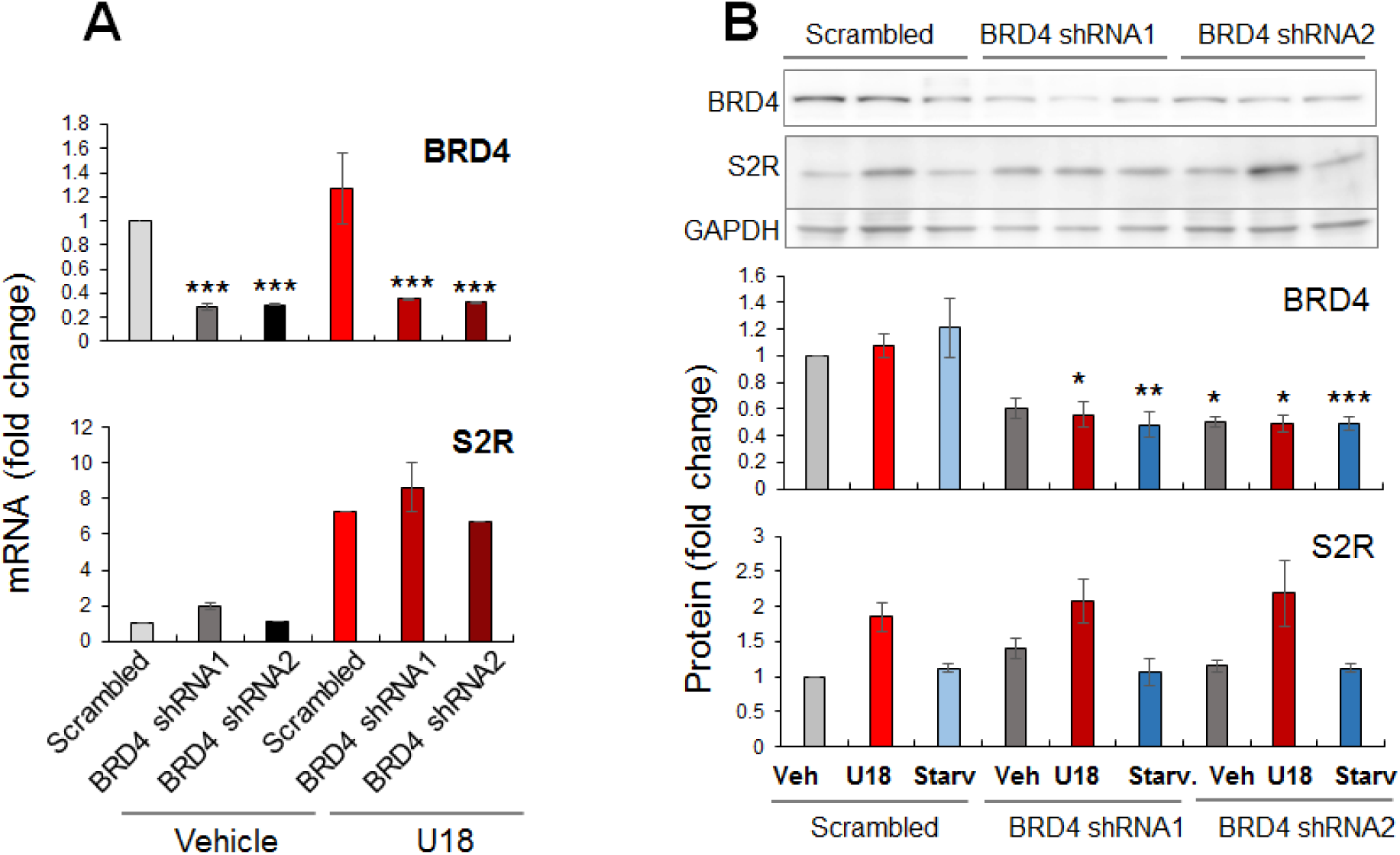
BRD4 silencing does not reduce S2R protein expression. Experiments were performed as described for Figure 2. Data presented here was obtained from the qRT-PCR (A) and Western blot (B) experiments using two shRNA sequences. The data indicates that experiments using both shRNAs led to similar results. Please refer to corresponding Figure 2C and 2F.

**Figure S3.**
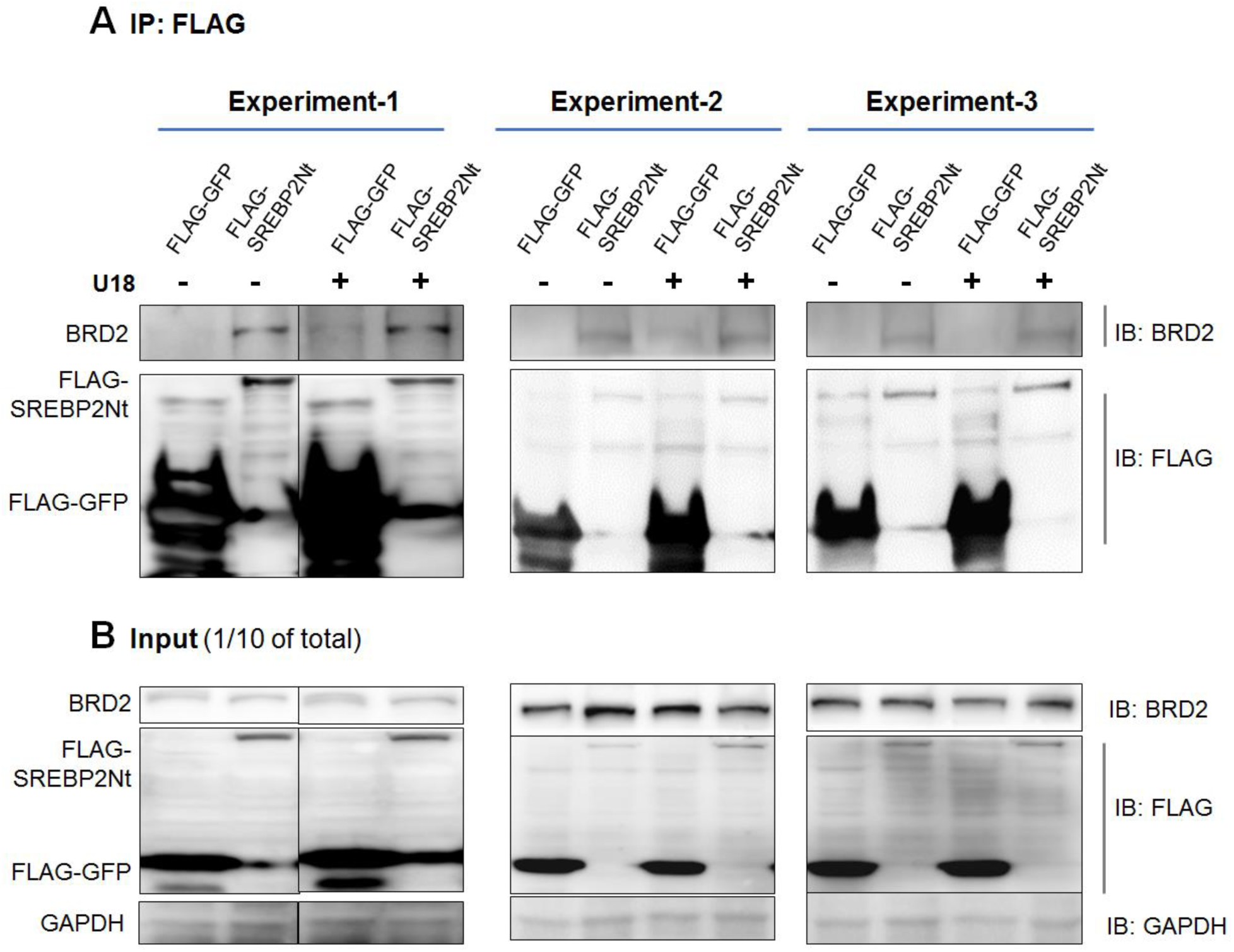
BRD2 co-IPs with the SREBP2 N-terminal TF half molecule. Experiments were performed as described in Figure 7A. Presented here are the full data from all three repeat experiments that produced the average values and bar graph in Figure 7A.

**Figure S4.**
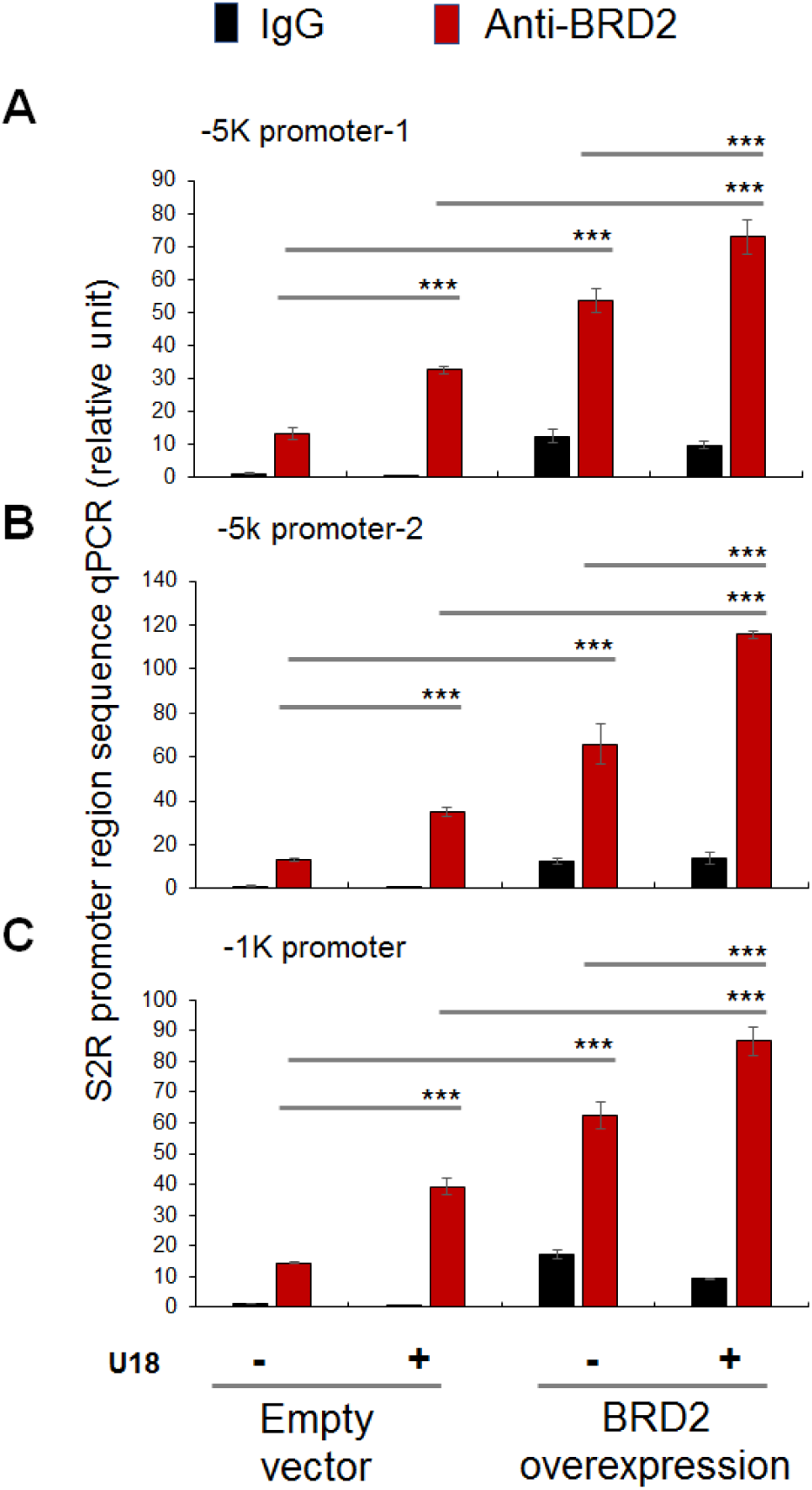
BRD2 occupies S2R gene promoter regions. ChIP-qPCR was performed as described in Figure 7B. Shown are results from all 3 experiments to detect BRD2 occupancy at 3 different S2R gene promoter regions, one of which (B) is presented in Figure 7B. The data indicate that all 3 experiments led to similar results.

**Figure S5.**
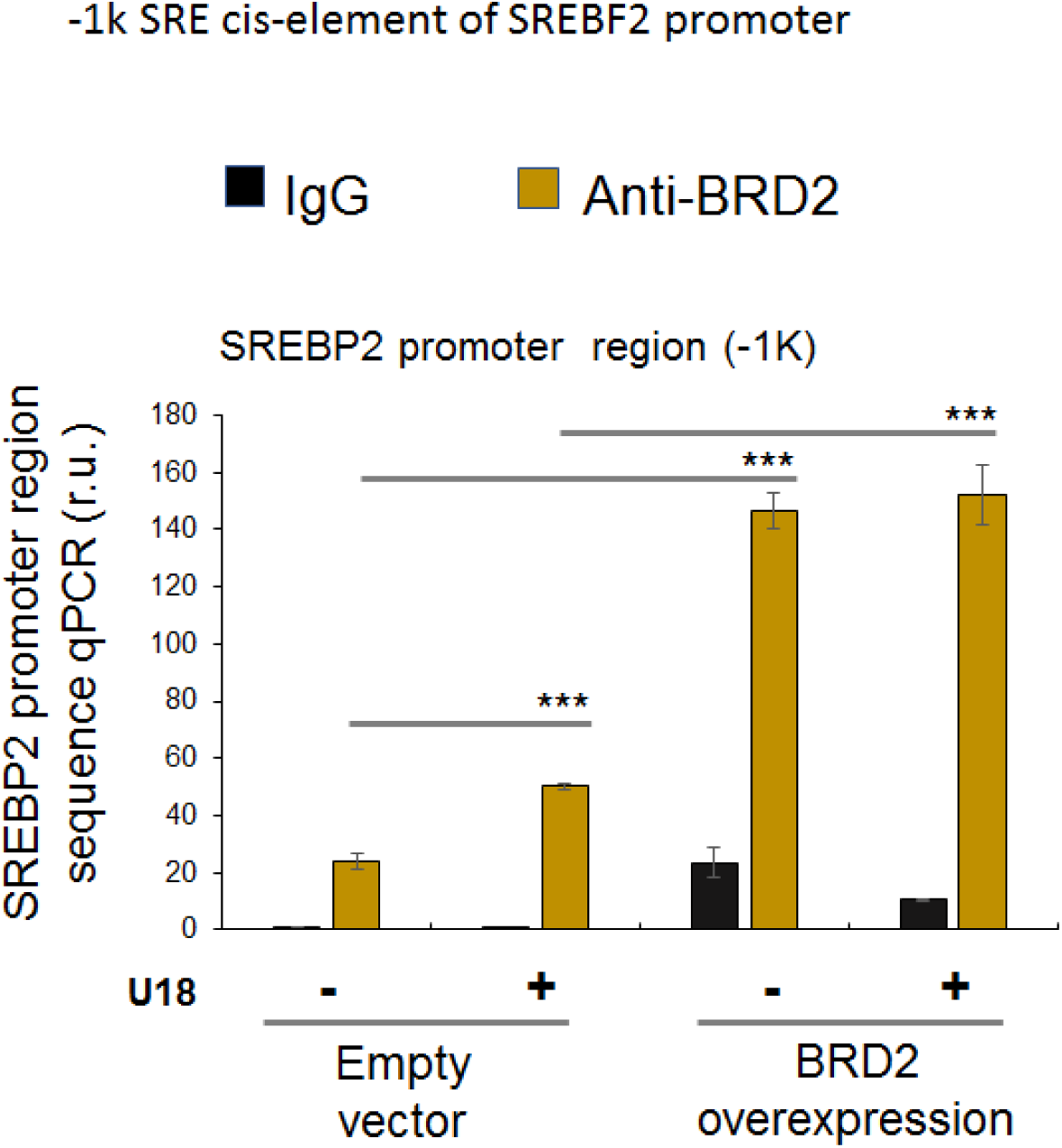
BRD2 antibody immunoprecipitates a SREBP2 gene promoter region. ChIP-qPCR was performed as described in Figure 7B. Shown is the experiment for the detection of BRD2 occupancy at the SREBP2 gene promoter. The data indicates that SREBP2 gene promoter (region) DNA was immunoprecipitated together with BRD2 by ChIP.

**Figure S6.**
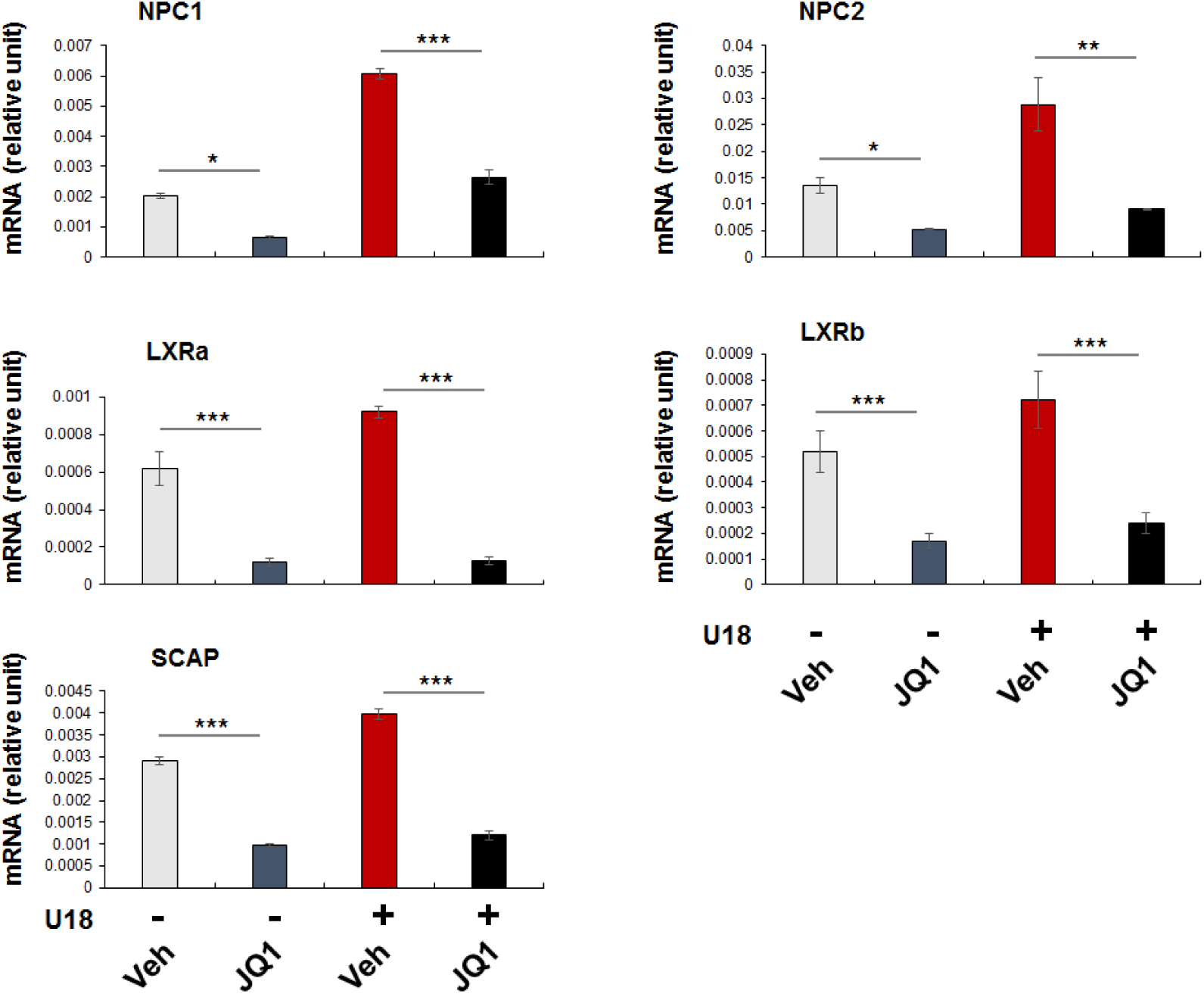
Pan-BETs inhibition attenuates the expression of SREBP target genes. Experiments were performed as described in Figure 1 except that several SREBP target gene mRNAs were assayed by qRT-PCR. Shown is each representative of two similar yet independent experiments. Statistics: Mean ± SD, n =3. One-way ANOVA with Bonferroni post-hoc test; *P<0.05, **P<0.01, ***P<0.001.

**Table S1.**
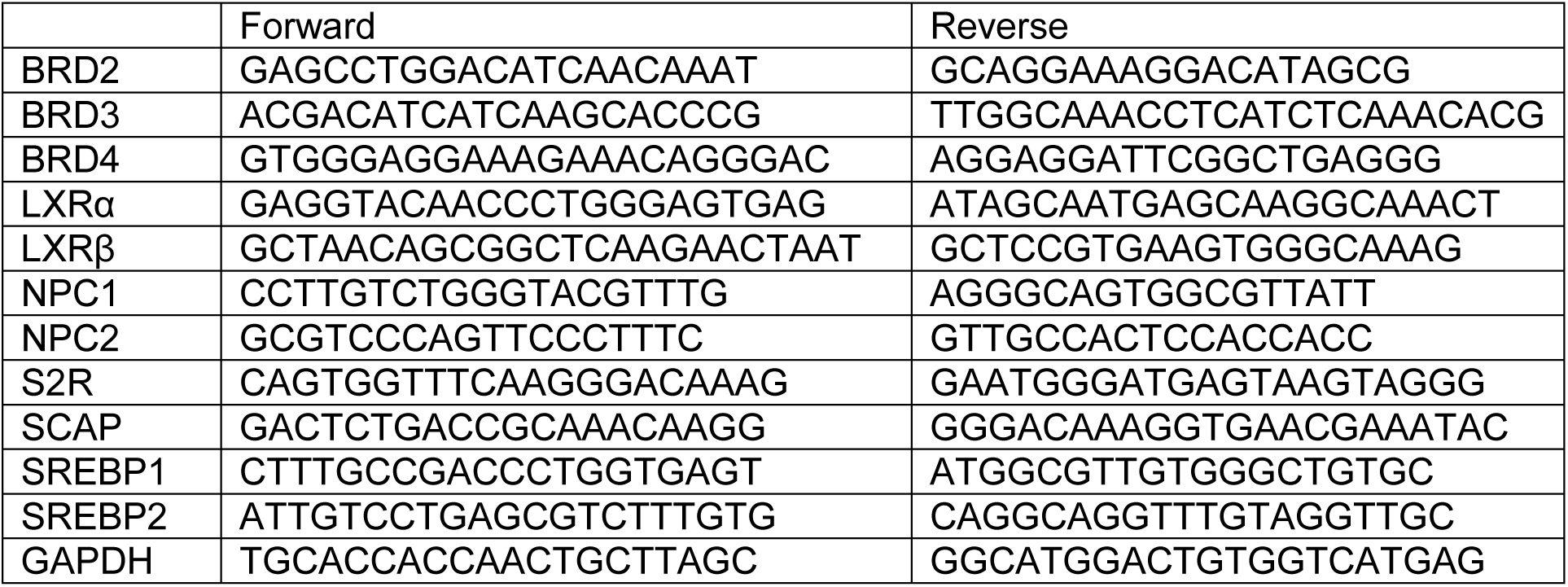
Primers used for qRT-PCR.

**Table S2.**
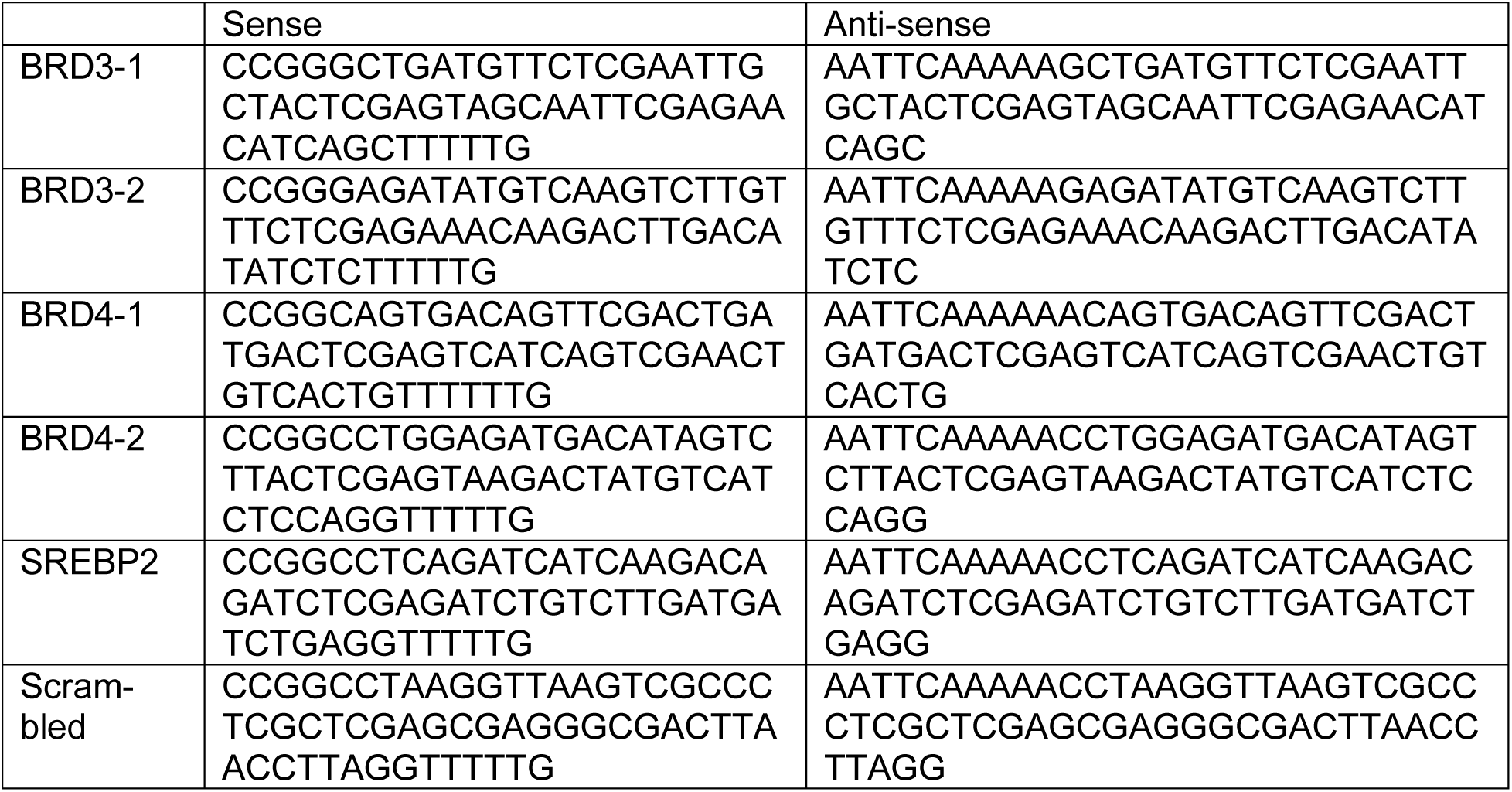
Oligonucleotide sequences for shRNAs.

**Table S3.**
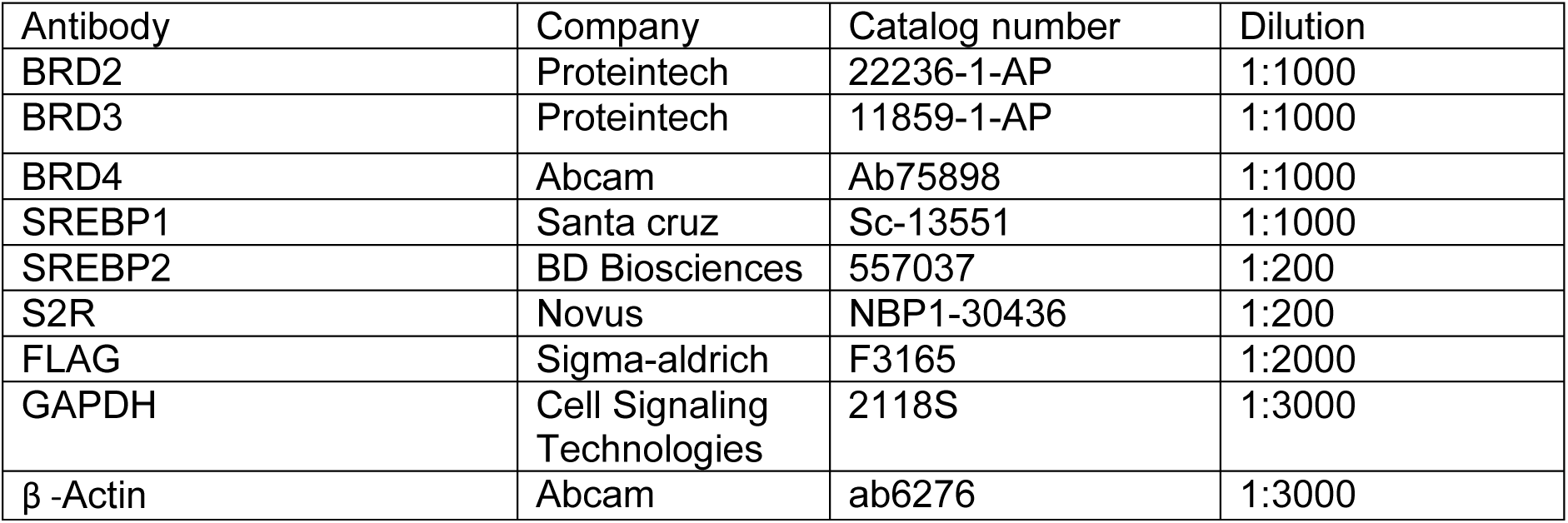
Antibodies for Western blotting.

**Table S4.**
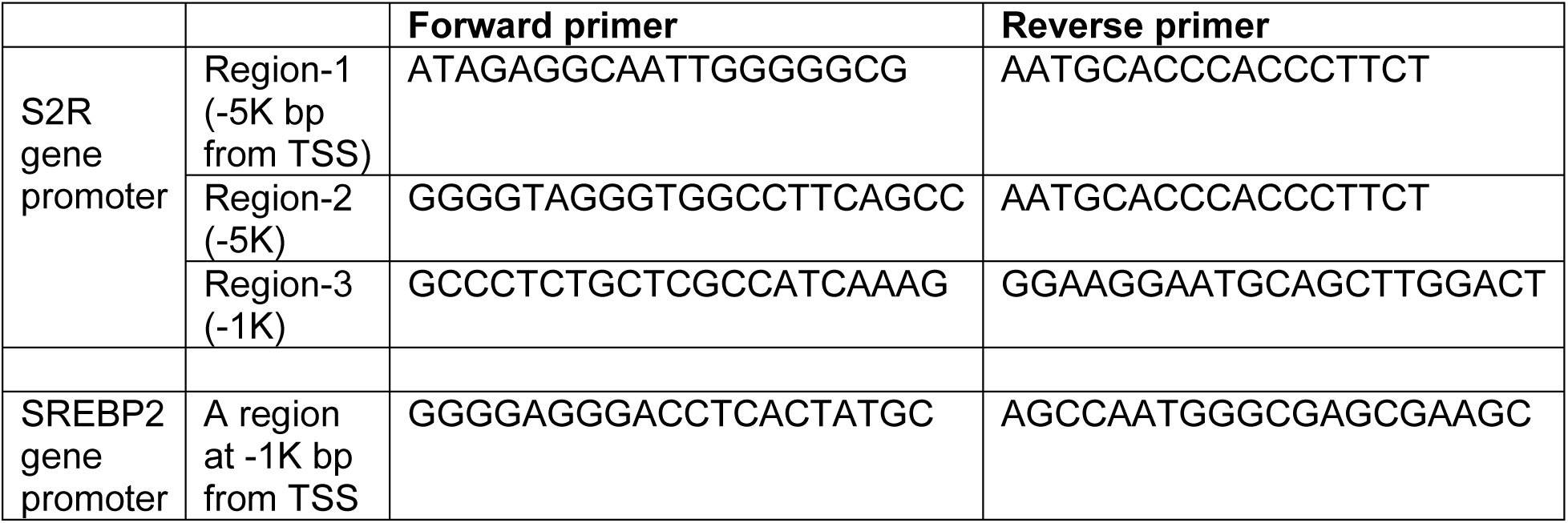
Primers for ChIP-qPCR.

